# Phased Chromosome-level Genome and Organellar Assemblies of *Castilleja foliolosa* Provide Vital Resource for Orobanchaceae Genomics

**DOI:** 10.64898/2026.09.21.753328

**Authors:** Ren N. Hamm, Jason Leung, Magdalene S. Lo, Adriana I. Hernandez, Sarah J. Jacobs

## Abstract

*Castilleja* (Orobanchaceae) is a diverse, facultatively hemiparasitic plant genus characterized by a complex evolutionary history of reticulate evolution and significant taxonomic ambiguity. High-quality genomic resources have historically been limited, hindering robust macro-evolutionary and macro-ecological inquiries. Here, we present the first high-quality, phased, chromosome-level reference genome for the diploid species *Castilleja foliolosa*. Utilizing a hybrid assembly strategy, we integrated long-read PacBio HiFi sequencing with Omni-C proximity ligation data to generate a highly contiguous nuclear assembly, complemented by reconstructed mitochondrial and chloroplastic genomes. The primary nuclear haplotype spans approximately 510 Mbp, containing 39,417 predicted genes and a substantial repetitive landscape covering ∼66% of the genome. Our organellar assemblies reveal notable structural complexity: the chloroplast maintains heteroplasmy via two primary structural haplotypes, while the 611 kbp mitogenome adopts a circular master topology existing in two isomeric forms. Comparative genomic analyses demonstrate a largely conserved chromosomal architecture relative to the closely related *Pedicularis cranolopha*, punctuated by the presence of lineage-specific genes. Functional enrichment of *Castilleja*-specific orthogroups identifies a consistent association with telomere maintenance, DNA integration, and zinc ion binding, likely stemming from the substantial repeat content. This reference genome provides a robust framework for dissecting the genomic drivers of taxonomic diversification and adaptation within the Orobanchaceae family. This resource significantly enhances our capacity to untangle reticulate evolutionary patterns, offering a definitive foundation for future comparative studies of genome evolution in the Orobanchaceae lineage of parasitic plants.

**Significance:** Modern conservation biology hinges on the assumption that species are discrete to one another, but actively radiating species complexes often maintain porous species boundaries and require high-quality genomic resources to effectively untangle dynamic taxonomic relationships and aid conservation assessments. In *Castilleja*, a clade characterized by reticulate evolution and ongoing gene flow across species boundaries, there is a growing need for high-quality reference genomes with greater utility for macro-evolutionary genetics. Here we present the first phased, chromosome-level *Castilleja foliolosa* reference genome as a resource suited for dissecting taxonomic uncertainty.

## Introduction

*Castilleja*, a widespread group of hemiparasitic wildflowers within the Orobanchaceae family known as the “paintbrushes,” exhibits a dynamic evolutionary history, hallmarked by repeated gene flow, hybridization, ploidy variability, and overlapping diagnostic morphological features resulting in many unresolved species complexes throughout the clade (Anderson & Taylor, 1983; Bürger & Chory, 2024; Heckard et al., 1980; Hersch-Green & Cronn, 2009; Jacobs et al., 2019; Tank & Olmstead, 2008, 2009; Wenzell et al. 2026, *in review*). Although they are primarily concentrated in western North America [Figure S1], these plants have undergone multiple hypothesized dispersals to South America (Tank & Olmstead, 2009), often experiencing cycles of introgression across species boundaries during recent glacial periods (Santos et al., 2026). Rapid diversification, coupled with morphological overlaps and wide ecological amplitude, has led to significant taxonomic ambiguity.

While genomic resources for within-genus studies have been gradually developed (Bürger et al., 2024, 2026; Latvis, Jacobs, et al., 2017; Latvis, Mortimer, et al., 2017), there remains a critical need for high-quality reference material to facilitate robust macro-evolutionary analyses, such as assessing heterozygosity, inferring demographic history, and investigating signals of selection and adaptation across the genus. To address these gaps, we focus our study on the diploid *Castilleja foliolosa* (commonly known as the Woolly Indian Paintbrush). *C. foliolosa* is a geographically widespread species, nested within the N. American southwest where much of the diversity of named species exists, and is closely related to multiple threatened species (Dunkle, 1943; Heckard et al., 1980). These concomitant factors highlight *C. foliolosa* as an excellent candidate for informing species conservation research, providing a foundational resource to better understand genome evolution within the genus and its broader family context.

## Results and Discussion

### Genome Assembly and Quality Assessment

This publication presents the first phased, long-read, annotated reference genome for *Castilleja*, encompassing the nuclear, mitochondrial, and chloroplastic assemblies. We first determined the ploidy of our *Castilleja foliolosa* individual using chromosome squash, confirming a diploid genetic makeup [Figure S2].

The *Castilleja foliolosa* genome was assembled by combining PacBio long-read and Omni-C short read data, resulting in a final HiRise assembly of 106 scaffolds with a GC content of 38.15% notable for a low frequency of sequence gaps (66, equating to ∼1.09 N per 100 kbp), and heterozygosity rate of 1.45% [Table 1]. Genome completeness was assessed using BUSCO scores against two databases: eukaryota (eukaryota_odb10) and embryophyta (embryophyta_odb10). This dual approach ensured that the results reflected both general genomic trends and specific plant gene diversity, respectively. In both cases, haplotype 2 maintained higher completeness and was selected for downstream annotation [Table 1].

**Table 1.**
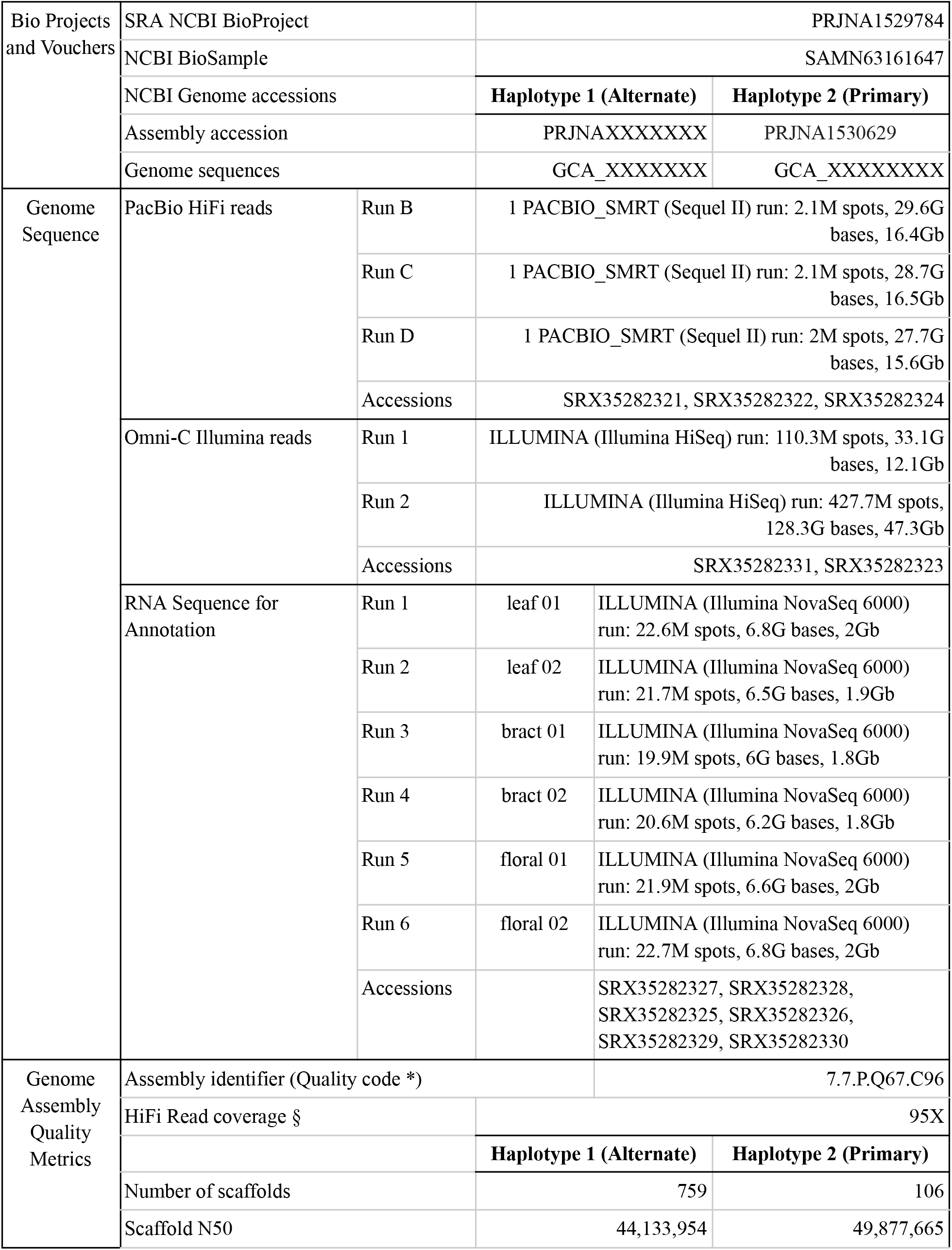
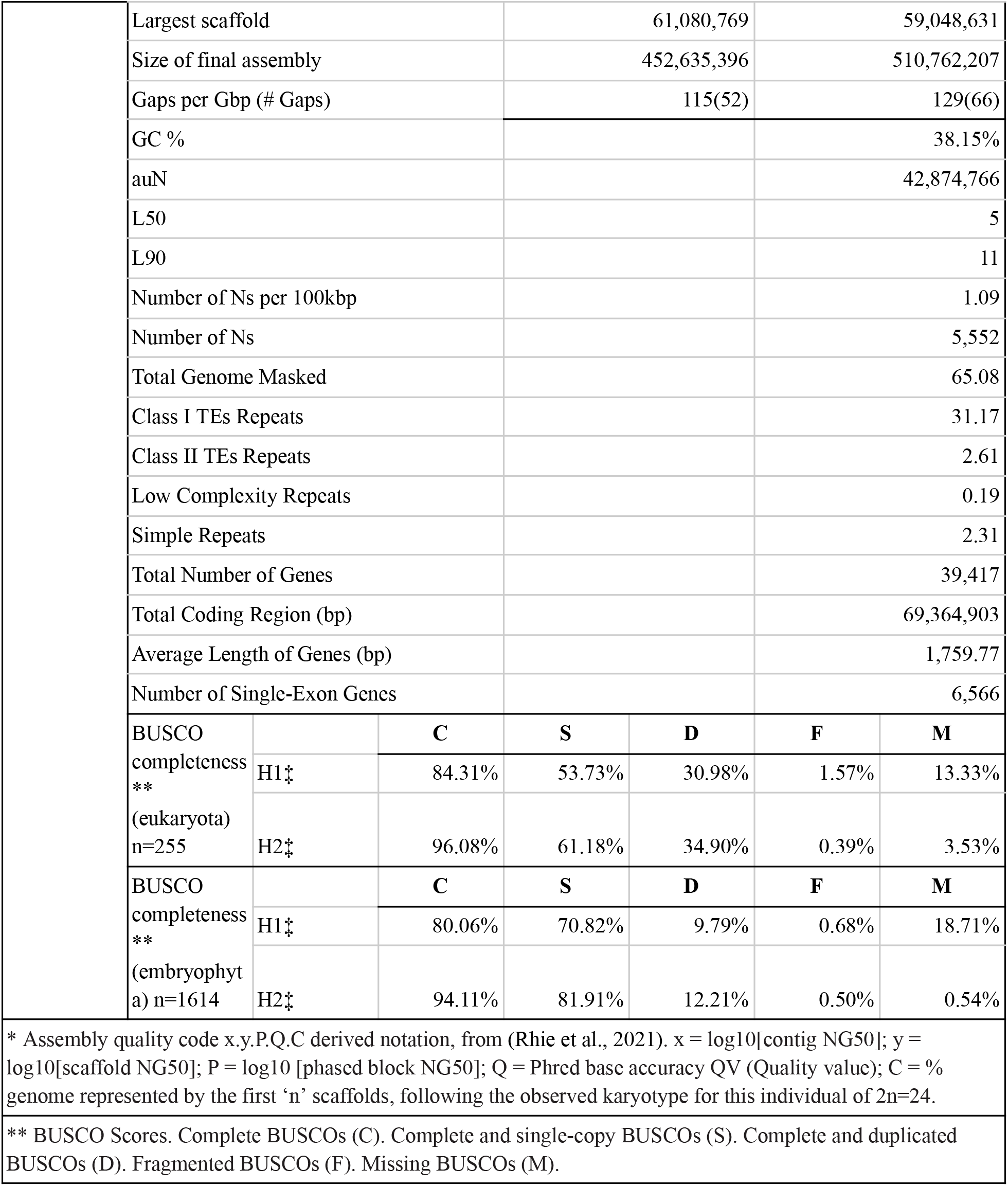
Nuclear genome assembly quality and contiguity statistics for *Castilleja foliolosa*. Alt text: Details include NCBI database accessions, raw sequencing run statistics across tissues (leaf, bract, floral), assembly structural features, repeat content, gene annotations, and BUSCO completeness estimates for two distinct haplotypes. Alt text: Details include NCBI database accessions, raw sequencing run statistics across tissues (leaf, bract, floral), assembly structural features, repeat content, gene annotations, and BUSCO completeness estimates for two distinct haplotypes.

Further annotation of the assembled sequence revealed several key features regarding the organism’s gene structure and content. The total number of genes predicted in the genome is 39,417, including 6,566 single-exon genes [Figure 1a]. The average genic length is 1,759.77 bp, and the total length of the coding regions is 69,364,903 bp. In addition to genic content, the repetitive content in the genome constitutes a total of 66.32% of the assembled sequence across 508,089 elements [Table S1; Figure 1a]. The most abundant class of repeats are LTR elements, which together account for 45.5% of the genome, with Gypsy repeats being the largest contributor at 18.79%. To quantify how much the identified transposable elements (TEs) in our genome have diverged over time from the sequences in the de-novo curated TE consensus library used for annotation, we examined the Kimura 2-parameter (K2P) distance scores calculated using the EDTA RepeatMasker option. As expected, the majority of these repeats minimally diverge, with ∼20% of the repetitive landscape diverging ≤1% from its representative consensus sequence [Figure S3]. Notably, we also observe a small percentage of TEs diverging up to 40% from their representative consensus sequence, indicating that these elements are older and have been subject to longer periods of mutational decay [Figure S3].

**Figure 1.**
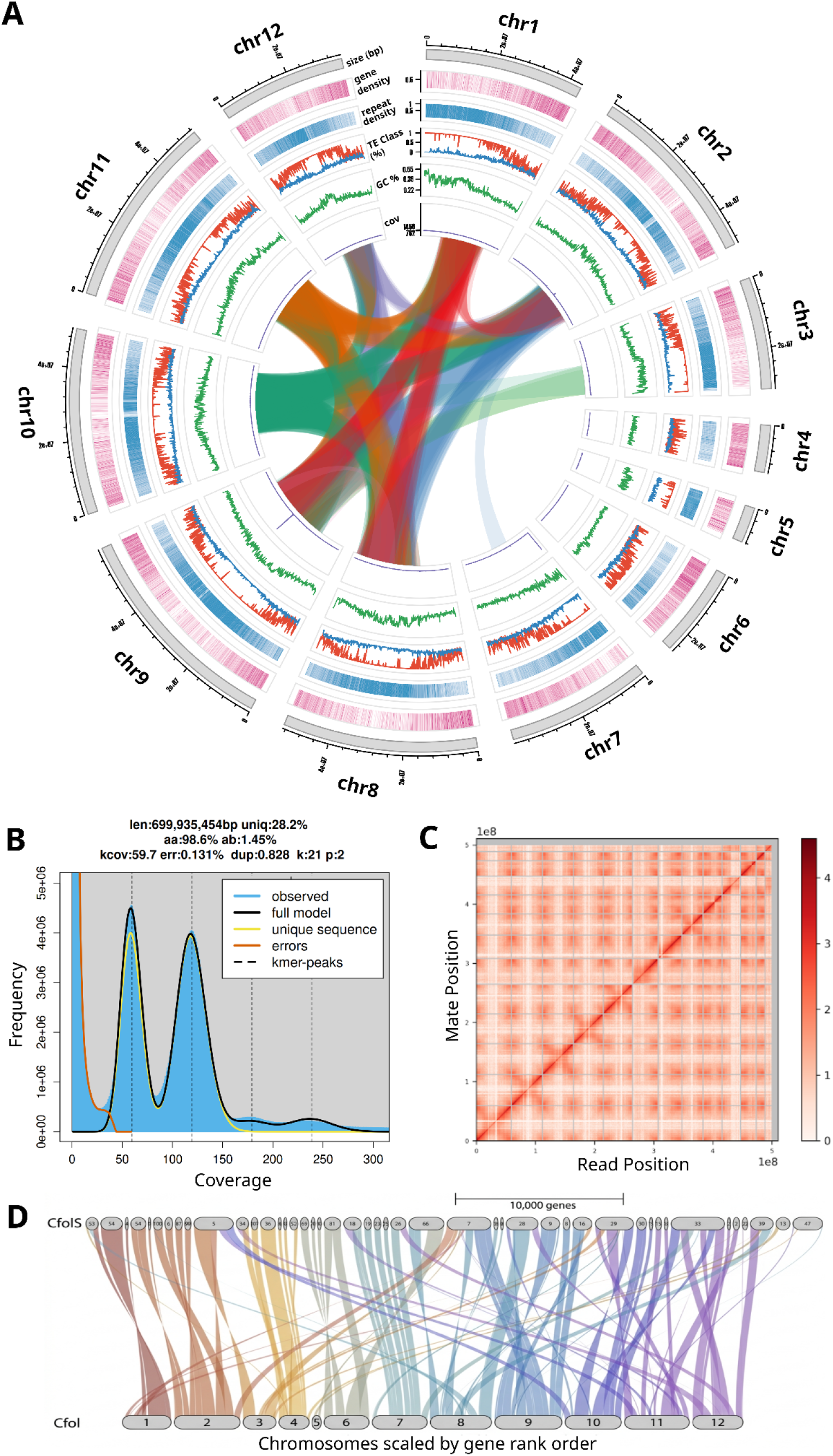
Architectural features, contact density, and comparative collinearity of the *C. foliolosa* genome assembly. A) Circos plot depicting nuclear genome assembly quality. Outer ring: chromosome length in megabases; Track 2: heatmap of gene density along the chromosome visualized in pink; Track 3: heatmap of total repeat density along the chromosome visualized in blue; Track 4: TE class identity as a ratio of total TE content along the genome, Class I retrotransposons are depicted in red and Class II DNA transposons are depicted in blue; Track 5: GC% visualized in green; Track 6: coverage % visualized in purple; Center: intra-genome syntenic collinearity passing a threshold of 150 genes consecutively across a >=5Mb span colored by chromosome. B) Hi-C heatmap displaying the normalized chromatin interaction frequencies across the 12 assembled chromosomes. The x- and y-axes represent the genomic position scaled in megabases, with black solid lines indicating chromosome boundaries. Interaction frequencies are binned at a 100-k resolution. The color intensity gradient corresponds to the log10-scaled contact density, where darker colors indicate a higher frequency of interactions. The prominent diagonal line demonstrates strong local intra-chromosomal interactions, confirming robust scaffolding and order of the reference assembly. C) Histogram showing the distribution of *k*=21-mer frequencies generated from raw PacBio HiFi sequencing data. The x-axis indicates *k*-mer coverage depth, and the y-axis represents the total count of unique *k*-mers at that depth. The empirical data density is indicated by the blue shading, while the red line tracks the optimal polyploid-aware mixture model fit calculated by GenomeScope 2.0. The bimodal peak distribution indicates a diploid genome. Inferred estimations for overall haploid genome size and heterozygosity rate are noted in the plot inset. D) The riparian alignment was generated using plot_riparian in GENESPACE, illustrating chromosome-scale gene order conservation. The newly presented *C. foliolosa* genome (bottom) and the previously published assembly (top) are stacked horizontally, with individual pseudo-chromosome and contig blocks, respectively, delineated along the axes. Colored ribbons represent highly conserved syntenic blocks connecting orthologous gene pairs across the two assemblies. Ribbon colors are mapped dynamically according to the chromosome IDs of the presented reference genome assembly. The x-axis chromosome segment scales are determined by gene rank order. Continuous, parallel ribbons indicate regions of strict structural conservation and collinearity, while crossing or passing ribbons highlight regions of structural variation, assembly orientation flips, or scaffolding refinements between the two genome versions. Alt text: A four-panel scientific figure labeled A through D illustrating the genome assemblies of the plant *Castilleja foliolosa*.Panel A is a multi-tracked circular Circos plot mapping nuclear genome features. Concentric rings display chromosome lengths, heatmaps for gene and repeat densities, transposable element class ratios, GC percentage, and read coverage, with a network of colored ribbons connecting syntenic regions across the center. Panel B is a square Hi-C chromatin interaction heatmap showing log10-scaled contact frequencies across twelve chromosomes, characterized by a sharp, prominent dark diagonal line and distinct grid-like boundaries. Panel C is a 21-mer frequency histogram displaying a bimodal peak distribution with blue empirical data shading and an overlapping red model fit line, including a text inset with genome size and heterozygosity metrics. Panel D is a linear, horizontally stacked riparian alignment plot comparing gene order between two genome versions, with thick, multi-colored ribbons connecting orthologous blocks across parallel horizontal axes.

The assembly reported here represents a significant improvement in quality and contiguity over the previously published *C. foliolosa* genome (Bürger et al., 2024), decreasing the 551 contigs to 106 scaffolds with 12 pseudo-chromosomes [Figure 1d], increasing the N50 value from 11,150,047 bp to 49,877,665 bp, and increasing the BUSCO completeness from 87.5% to 96.1% [Table S2].

### Chloroplast Genome Maintains Structural Heteroplasmy

The chloroplast assembly for *C. foliolosa* revealed the presence of at least two predominant structural cp haplotypes [Figure S4a]. While long reads enabled the *de novo* assembly of contigs that collectively span the entire expected chloroplast genome, the observed haplotypes prevented the final extraction of a single contig [Table S3], and indicate substantial structural heteroplasmy. The manually assembled genomes for cpHaplotype 1 and cpHaplotype 2 were identical in length, totaling 152,953 bp each, but differed in the order and direction of their quadripartite structure lengths (LSC, SSC, and IR regions) [Figure S4a]. By subsequently quantifying long read coverage of all possible structural haplotype orientations, generated from the identified LSC, IRs, and SSC by including complements and reverse complements, we confirm the presence of only two major haplotypes. The vast majority of reads align to Cp-hap assemblies where the LSC and IR are adjoined as is. Reads spanning the IR1-SSC junction show an IR_SSC to IR_SSCrc (reverse complement) ratio of approximately 2:1, and reads spanning the IR2-SSC junction all map to the reverse complement of IR2, suggesting only two structural haplotypes, congruent with established models of plastid structural dynamics in angiosperms (Lee et al., 2020; Witharana et al., 2026).

### Mitochondrial Genome Maintains Two Isomeric Topologies

The *C. foliolosa* mitogenome was assembled into a 611,107 bp circular master topology [Figure S4b, S5, S6]. To prevent coverage inflation from plastid-derived Mitochondrial Plastid DNA (MTPT), a dual-record reference mapping approach was deployed, resolving single-reference depth spikes and establishing a uniform canonical mitochondrial coverage profile (mean: 3,495×, median: 3,616×) [Figure S7]. Annotation via PGMA identified 74 total loci, which were intersected with 30 identified MTPT regions to differentiate genuine mitogenic genes from plastid insertions [Figure S5]. An additional 10 genes shared plastid homology, while 11 fully MTPT-resident loci were flagged as plastid-derived pseudogenes [Figures S4, S5, S6]. GeSeq annotation localized the 30 MTPT regions across ∼79 kb (∼13% of the mitogenome), tracing 29 regions back to specific chloroplast donor genes involved in photosystems, ATP synthase, and RubisCO [Table S4].

Structurally, the master circle comprises three primary segments (u33, u34, and u35). Read overlap analysis confirmed a circularized junction between u34 and u35, while a 37 bp inverted repeat at the u33–u34 boundary mediates frequent in vivo recombination. This mechanism drives a structural inversion of the u34 segment, maintaining two co-existing isomeric topologies within the plant population: the u34+ orientation at ∼54% abundance and the u34− orientation at ∼46% abundance [Table S6, S7].

### Karyotype Diversification in C. foliolosa

To examine the evolutionary dynamics of chromosomal rearrangements and gene content diversification within this lineage, we compared our genome with the closely related, high-quality *Pedicularis cranolopha* reference genome. Syntenic depth analysis using MCScanX revealed a predominant 1:1 correspondence between *P. cranolopha* and *C. foliolosa* [Figure S8a]. Approximately 25-30% of genes in each genome were present in single-copy syntenic blocks, with only a small fraction (9%) showing depth-2 relationships, indicating the absence of recent whole-genome duplication and supporting largely conserved chromosomal architecture [Figure S8a]. We also observe evidence of alternative genomic structural variation, including a large inversion on *C. foliolosa* chromosome 12 corresponding to *P. cranolopha* chromosome 6 [Figure S8b,c], and breaks in some synteny diagonals, possibly due to centromere transposon content [Figure S8b,c].

A large proportion of *C. foliolosa* genes identified using Orthofinder lack detectable synteny (approximately 61%) across 10 reference species [Table S7], indicating a substantial number of lineage-specific genes. Notably, of the 36,384 *C. foliolosa* orthogroups, 1,025 are only found within the *Castilleja* lineage, encompassing 22.8% of the genes found. Functional enrichment analysis of *Castilleja*-specific orthogroups reveals a consistent association with telomere maintenance (GO:0000723, p= 2.55E-25) and DNA integration (GO:0015074, p=1.69E-21) confirming the prevalence of retrotransposons [Table S8]. Notably, the significantly enriched GOterm related to the highest number of *Castilleja*-specific genes relates to Zinc Ion Binding (GO:0008270, p=1.69E-21). Given that many hemiparasitic species actively accumulate metal ions from their hosts, this enrichment may reflect specific evolutionary pressures on nutrient absorption networks or stress defenses active at the haustorial interface (Adler, 2002; Cabot et al., 2019).

## Materials and Methods

### Sample Collection and Description

We collected tissue from a single individual of *Castilleja foliolosa* in February 2022 near Northern California’s Putah Creek. For DNA and RNA sequencing, we collected young leaf and floral tissue flash frozen in liquid nitrogen immediately after collection to prevent DNA degradation.

### Chromosome Squash

Chromosome squashes were performed following the general protocol of (Windham et al., 2020) with modifications optimized for *Castilleja* floral buds.

### Sample Preparation and Sequencing

DNA was extracted using a modified CTAB (Doyle CIT) protocol. DNA samples were quantified using Qubit 2.0 Fluorometer (Life Technologies, Carlsbad, CA, USA). The PacBio SMRTbell library (∼20kb) for PacBio Sequel was constructed using SMRTbell Express Template Prep Kit 2.0 (PacBio, Menlo Park, CA, USA) and bound to polymerase using the Sequel II Binding Kit 2.0 (PacBio). Sequencing was performed on PacBio Sequel II 8M SMRT cells. Total RNA was extracted from leaf and floral tissue, quantified via Qubit 2.0, and quality-asserved via Agilent 2100 Bioanalyzer. Stranded libraries were prepared using the Illumina Stranded mRNA Prep Kit via magnetic oligo(dT) poly(A) selection and sequenced on an Illumina NovaSeq 6000 platform to generate paired-end 150 bp reads for structural genome annotation.

### Nuclear Genome Assembly

85.8 Gb of PacBio CCS reads and 133 Gb of OmniC data were used as an input to HiC integrated hifiasm with default parameters. Blast results of the output assemblies against the nt database were used as input for blobtools2 v1.1.1 (Laetsch & Blaxter, 2017), and scaffolds identified as possible contamination were removed from the assembly. Finally, purge_dups3 v1.2.5 (Guan et al., 2020) was used to remove haplotigs and contig overlaps. Omni-C sequencing libraries produced by Dovetail (Scotts Valley, CA, USA) were generated using NEBNext Ultra enzymes and Illumina-compatible adapters. Libraries were sequenced on an Illumina HiSeqX platform to produce a approximately 30x sequence coverage. Then HiRise used MQ>50 reads for scaffolding [Figure S9]. HiRise (Putnam et al., 2016) was used to scaffold the de novo assemblies with OmniC library reads. Following alignment to the draft assembly using bwa (https://github.com/lh3/bwa), HiRise modeled genomic distances from read pair separations to identify, break, and correct misjoins.

### Chloroplast Genome Assembly

Long reads were mapped to the *Castilleja paramensis* chloroplast genome (Fan et al., 2016) with minimap2 v.2.24 (Li, 2018) and aligned to an altered version of the reference with the halves flipped to ensure that reads which spanned ends of the original were not excluded. To assemble the reads, we used Canu v.2.2 (Nurk et al., 2020) with the “-pacbio-hifi” option and a genome size estimate of 160,000 bp.

The two largest contigs were initially annotated with GeSeq (Tillich et al., 2017) to ensure adequate coverage when compared to *C. paramensis* and used to divide the contigs at the boundaries of the long single copy region (LSC), short single copy region (SSC), and inverted repeats (IRa and IRb). We employed Cp-hap (Wang & Lanfear, 2019) to identify the structural haplotypes within our reference individual using realignments of all long reads to different combinations of the quadrapartite regions and their reverse complements. After ensuring correct orientation, GeSeq was used for final annotation, and OGDRAW (Greiner et al., 2019) was used for visualization.

### Mitochondrial Genome Assembly

We assembled the *C. foliolosa* mitochondrial genome using oatk v1.0 (PMID: 40775726), conducted with the *embryophyta* database and a minimum k-mer coverage of 500, yielding a unitig assembly graph (GFA) from which a master path consensus sequence of 611,107 bp was derived.

To validate the closure and identify plastid-derived DNA (MTPT), we constructed a dual-record reference including the assembled mitochondrial sequence appended with its first 25 kb to facilitate mapping wrap-spanning reads and previously assembled plastid sequence. HiFi reads were aligned using minimap2 (*-ax hifi --sam-hit-only --secondary=no*) to ensure that plastid-derived reads preferentially mapped to the plastid record. Further read mapping was conducted using the mitochondrial sequence alone as a reference to identify MTPT sections. Potential structural isoforms were identified via links and sequence overlaps within the GFA file. Relative isoform frequency was estimated using both the EC counts generated by oatk and the read mapping data utilized for circularity and MTPT verification.

Functional annotation was performed using the PGMA web server, employing tRNAscan-SE (Lowe & Eddy, 1997) for the tRNA identification and OGDRAW (Greiner et al., 2019) for visualization.

### Haplotype Quality Assessment and Selection

We assessed haplotype completeness using Benchmarking Universal Single-Copy Orthologs (BUSCO) v5.4.4 (Simão et al., 2015) with the eukaryota_odb10 (species: 70, BUSCOs: 255) and embryophyta_odb10 databases (species: 50, BUSCOs: 1,614). Haplotype 2 had the highest completeness across both methods and was chosen for downstream annotation. To estimate heterozygosity, raw reads were profiled using GenomeScope 2.0 based on a 21-mer frequency distribution generated by Jellyfish (v2.2.3).

### Genome Annotation

Repeats were identified de novo using RepeatModeler (version 2.0.1; Flynn et al., 2020) and masked using RepeatMasker (Version 4.1.0). Transposable elements were identified de novo using EDTA (Extensive de-novo TE Annotator, v1.10.9; Ou et al., 2019) with default parameters.

Coding sequences from *Arabidopsis thaliana* (GCA_020911765.2), *Buddleja alternifolia* (GCA_019426215.1), *Paulownia fortunei* (GCA_019321725.1), *Salvia splendens* (GCA_020531475.1), *Sesamum indicum* (GCF_000512975.1), and *Solanum lycopersicum* (GCA_000188115.5) were used to train two ab initio models for *Castilleja foliolosa*: one using AUGUSTUS software (version 2.5.5; Stanke et al., 2008) with six rounds of optimisation and another using SNAP (version 2006-07-28; Korf, 2004). RNAseq reads were mapped using STAR (Version 2.7), and intron hints were generated using bam2hints tools within AUGUSTUS. MAKER, SNAP and AUGUSTUS were then used to predict genes in the repeat-masked reference genome. Swiss-Prot peptide sequences were used in conjunction with protein sequences from the training species to generate peptide evidence in the Maker pipeline. Only genes predicted by both SNAP and AUGUSTUS softwares were retained in the final gene sets, and AED scores were generated for each predicted gene. Genes were further characterised for putative function via BLAST search of the peptide sequences against the UniProt database. tRNAs were predicted using the software tRNAscan-SE version 2.05; (Lowe & Eddy, 1997).

### Comparative Genomics

We compared our reference genome to six species in the Orobanchaceae: *Lindenbergia luchunensis, Striga asiatica, Phtheirospermum japonicum, Pedicularis cranolopha, Phelipanche aegyptiaca, Orobanche cumana*, and three outgroup species (*Coffea canephora, Mimulus guttatus, Olea europaea*) utilized in previous family-level comparative genomics research [Table B] (Xu et al., 2022a). Amino acid sequences of all 10 combined reference genomes’ gene models were used as input into GENESPACE (Lovell et al., 2022) to run Orthofinder (Version 2.5.5; Emms & Kelly, 2019), using DIAMOND (Version 2.1.7; Buchfink et al., 2021) for local sequence alignment. mRNA sequences were used to estimate each whole genetic feature from our gff files and CDS fields to specifically investigate conserved regions more representative of evolutionary change.

We further examined the 1,025 *Castilleja*-specific orthogroups by assessing functional enrichment using Gene Ontology (GO) terms inferred through sequence homology (Thomas et al., 2022). Because *Castilleja* lacks curated GO annotations, functional inference was performed by transferring GO terms from the closely related *Mimulus guttatus* (IM62 v3.1; Lovell et al., 2025) reference genome using BLASTP implemented in Geneious Prime (Version 20205.2; https://www.geneious.com) retaining the top-scoring hit per query above an e-value threshold of ≤ 1e-05. GO annotations were retrieved using Phytozome’s Biomart Tool (Goodstein et al., 2012). Enrichment was performed using Planteome’s Ontology Enrichment Analysis Tool (https://planteome.org/oat/), applying a chi-squared test.

Using the above GENESPACE analytical framework, we additionally evaluated syntenic conservation between the assembly generated in this study and the previously published assembly (Bürger et al., 2024).

## Data Availability

The raw sequencing data, assembled genome, and organelle sequences have been deposited in the NCBI Sequence Read Archive (SRA) under BioProject accession PRJNA1529784.

Primary and alternate genome assemblies are available under BioProject accession numbers PRJNA1530629 and PRJNXXXXXX, respectively. Supporting transcriptome datasets are available under BioProject PRJNA1529784.

## Acknowledgements

We thank Jim Henderson for his diligent work assembling the mitochondria, utilizing the resources provided by the California Academy of Sciences’ Center for Comparative Genomics. This work was supported by Dovetail’s Tree of Life Award [to SJJacobs]; and the Rose Postdoctoral Fellowship, both of which provided financial support essential to the success of this project.

## Notes

### Competing Interest Statement

The authors have declared no competing interest.

