## Supplemental Figures S1-S9 and Tables S1-S8 for "Phased Chromosome-level Genome and Organellar Assemblies of *Castilleja foliolosa* Provide Vital Resource for Orobanchaceae Genomics"

### Supplementary Figures:

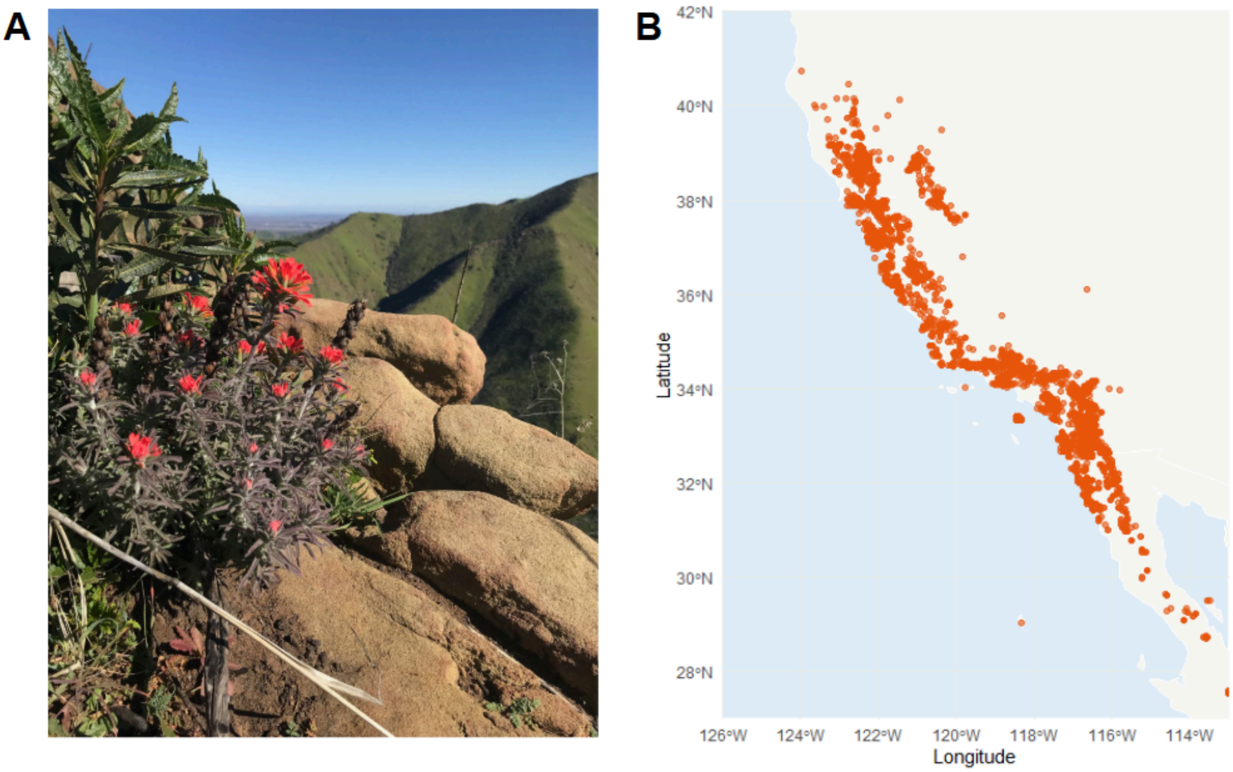


**FIGURE S1: Phenotype and Range of *Castilleja foliolosa*.** A) Sequenced individual used for genome assembly. Photo credit: Sarah Jacobs, February 1st 2022. B) Distribution map of all known occurrences of *Castilleja foliolosa* on the GBIF database. [GBIF.org (18 September 2026) GBIF Occurrence Download https://doi.org/10.15468/dl.yw7k6g].

Alt text: Photograph taken of the sequenced individual at the field location paired with a distribution map of the known range of the species.

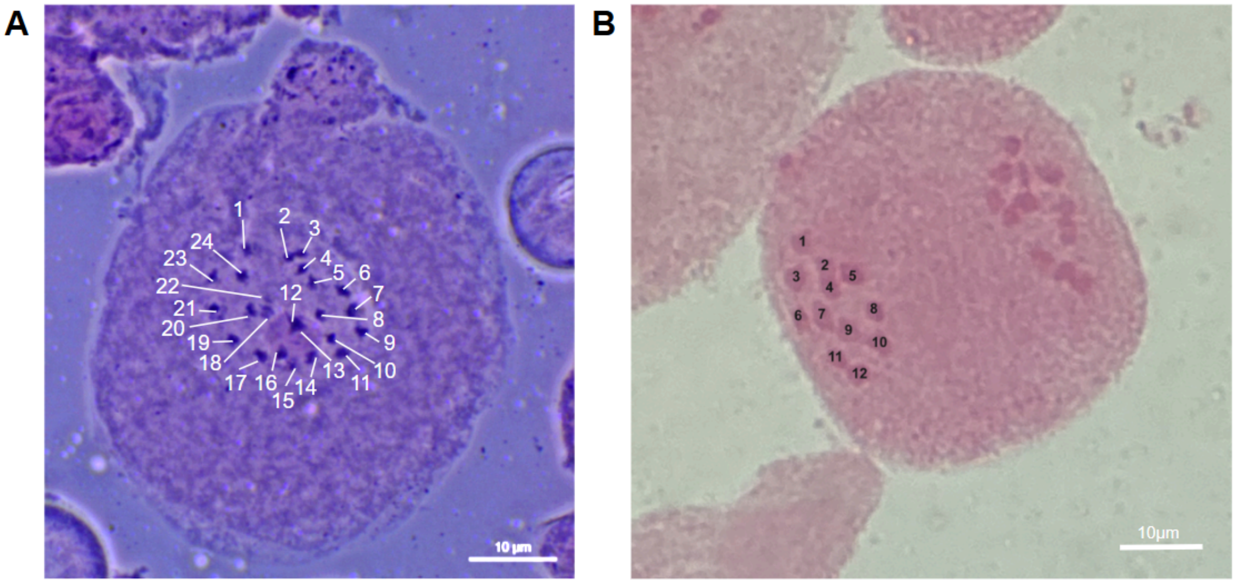


**FIGURE S2: Castilleja foliolosa individual confirmed as diploid.** A) Chromosome squash of developing floral bud cell in prophase from the sequenced individual. Diploid chromosome counts are depicted in white numbers totalling 2n = 24. B) Chromosome squash of developing floral bud cell in anaphase from a neighboring individual approximately 3 feet away in distance. Chromosome counts are depicted in black numbers totally 1n = 12.

Alt text: Two photos side by side of chromosome counts from chromosome squashes. The photo on the left enumerates all 24 chromosomes in the sequenced diploid individual. The photo on the right enumerates 12 chromosomes on one side of the dividing cell from a neighboring individual to establish consistency across the population.

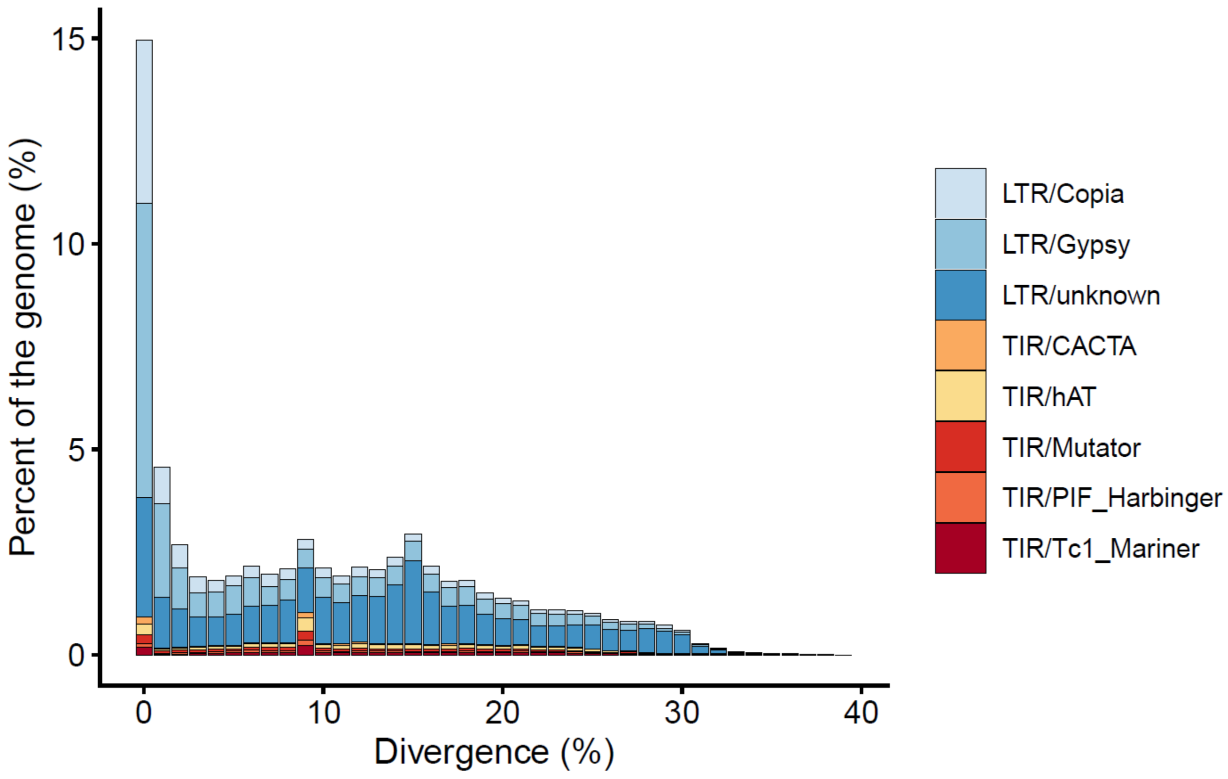


**Figure S3: Transposable element divergence highlights complex history of transposable element activity.** The distribution of sequence divergence (Kimura 2-parameter distance) across identified transposable element (TE) families relative to the *de novo* curated consensus library generated from our genome. Lower divergence scores indicate recent transposition events, while higher divergence reflects mutational decay over longer evolutionary timeframes. Abbreviations: LTR = long terminal repeat, TIR = terminal inverted repeat.

Alt text: Stacked bar chart counting the distribution of sequence divergence (Kimura 2-parameter distance) in transposable elements (TEs) across the genome. The majority of TEs exhibit divergence around ~0%, but TEs covering ~3% of the genome exhibit divergence estimates up to 30%, highlighting activity across large timescales.

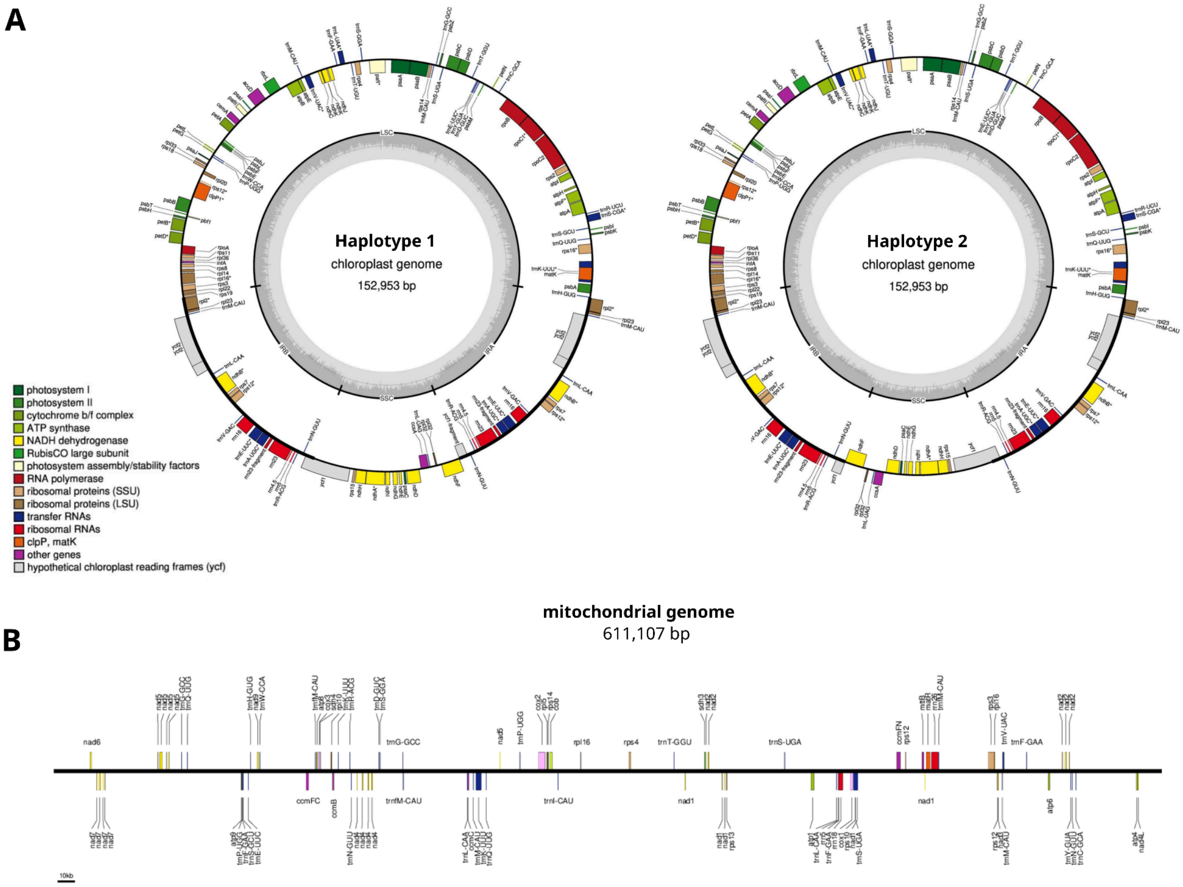


**Figure S4: Annotated Organellar Assemblies.** A) The circular diagrams represent the two distinct, co-existing structural orientations (Haplotype 1: LSC-IRa-SSC-IRb; Haplotype 2: LSC-IRa-SSCrc-IRb) isolated from the assembly, differing primarily by the inversion of the Small Single-Copy (SSC) region. Both genomes exhibit a characteristic quadripartite architecture, containing a Large Single-Copy (LSC) region, a Small Single-Copy (SSC) region, and a pair of identical Inverted Repeats (IRa) and (IRb). Genes drawn on the inside of the outer circle are transcribed in the clockwise direction, while those on the outside are transcribed counterclockwise. Genes are color-coded into standard functional groups as specified by the accompanying color index. The inner gray circle outlines the overall GC content across the plastome sequence. B) The physical map displays the complete mitogenome sequence, linearized starting at the nad6 gene for standardized visualization. Genes positioned above the central coordinate bar are encoded on the forward strand and transcribed from left to right, whereas genes below the bar are encoded on the reverse strand and transcribed from right to left. Features are color-coded according to functional groups following standard OGDRAW functional nomenclature.

Alt text: Panel A shows two circular plastid genome diagrams with a classic quadripartite architecture. Panel B is a linear physical map of a mitochondrial genome, displaying functional gene blocks color-coded by category above and below a central coordinate timeline starting at the *nad6* gene.

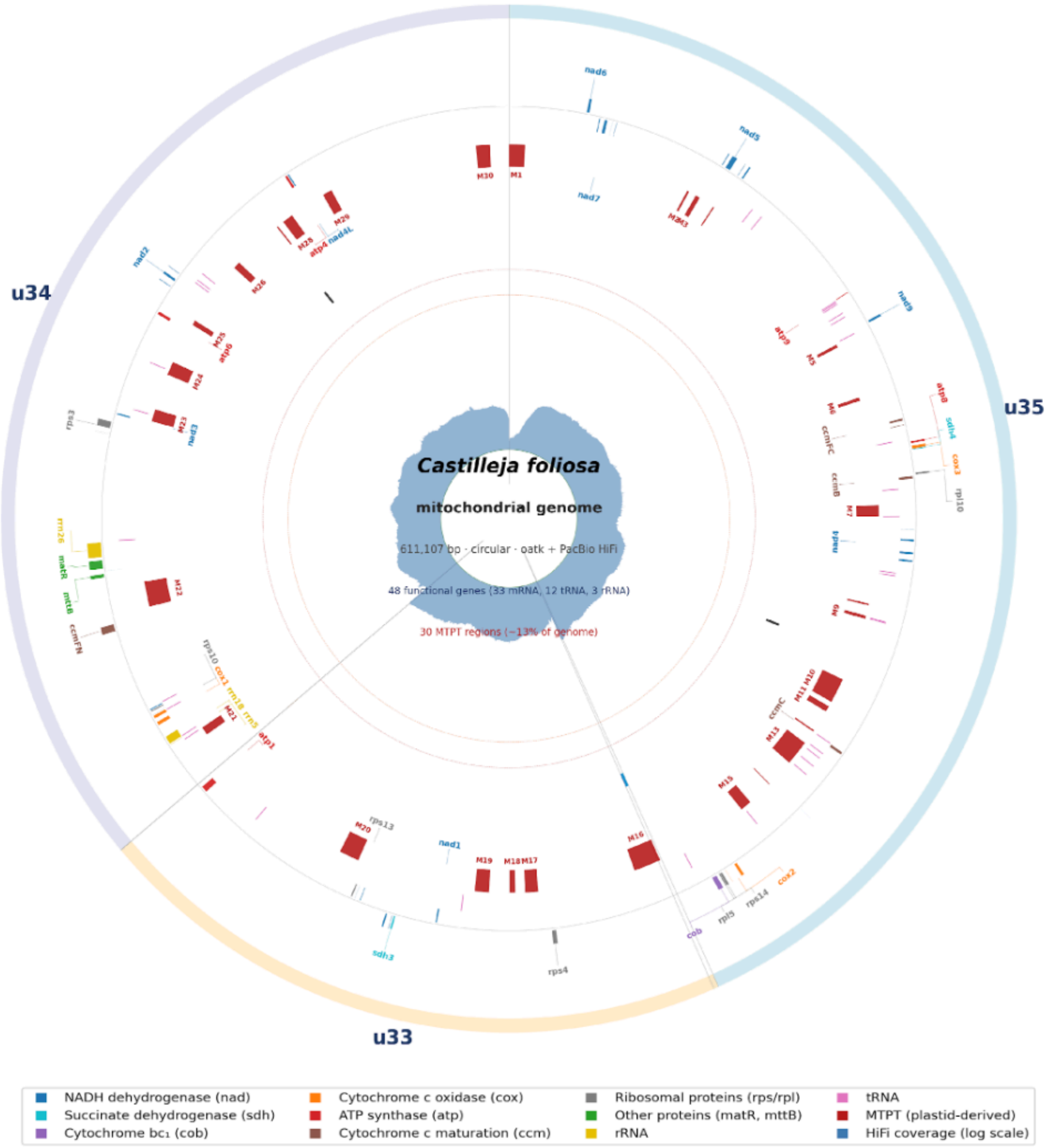


**Figure S5. Mitochondrial Genes, MTPT, and Coverage.** Integrated circular map of the 611,107 bp Castilleja foliosa mitochondrial genome. Tracks (outer to inner): contig identity ring (u35, u33, u34) on the master path; mitochondrial gene exons split by strand (+ strand outer band, − strand inner band) and colored by functional category (gene-name labels with leader lines on both sides); tRNA tick marks; the 30 MTPT regions (red blocks, M1–M30 labeled); the four recombination repeats discussed in the isoform analysis (the 958 bp u33↔u35 direct repeat at the u33/u35 boundary, the 37 bp u33↔u34 inverted repeat at the u33/u34 boundary, the 44 bp u35 self-loop direct repeat, and the new 740 bp / 87% identity inverted repeat between u34 and u35 shown as two paired markers); and innermost — per-base HiFi coverage on a log scale (light blue fill). The coverage track shown here is from the plastid mapping (HiFi reads aligned to a custom reference of the mitochondrion plus the chloroplast as a separate record), which maps majority of MTPT-derived plastid reads onto the plastid record and reveals a near-uniform ~3,000–3,600× canonical mitochondrion coverage. The dotted reference ring at ~3,200× marks the mitochondrion baseline.

Alt text: Circular map of mitochondrial assembly divided into three contig identities and annotated for genes colored by known function. Coverage is depicted as a wrapping bar chart in the center.

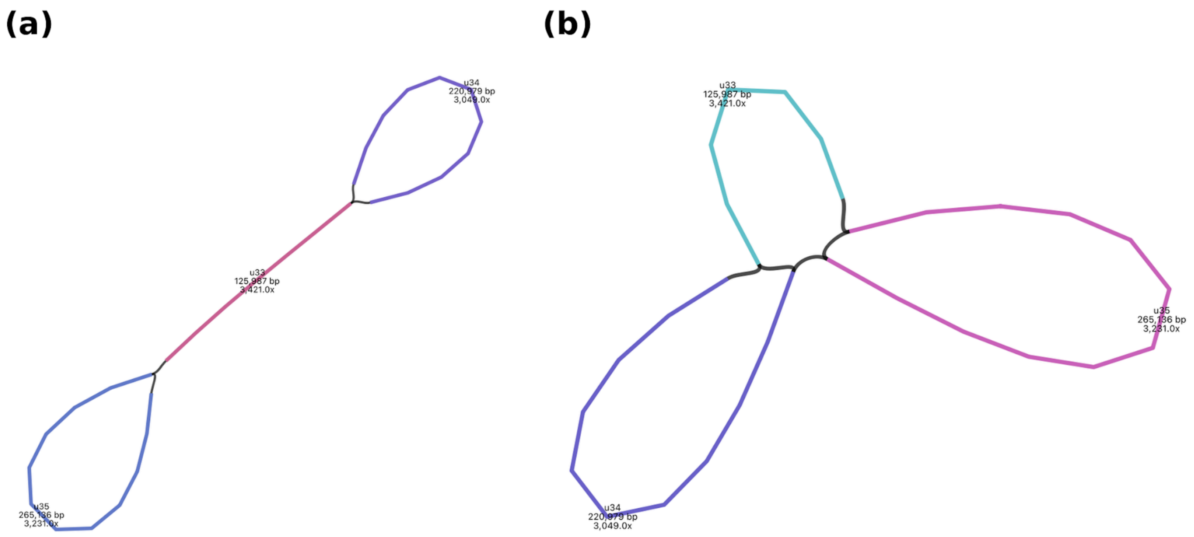


**Figure S6: Barbell and Master Circle Graphs.** BandageNG rendering of the C. foliosa mitochondrial assembly graph before and after adding a master-circle closure L-record. A) Original oatk-generated assembly graph with three unitigs — u33 (125,987 bp), u34 (top loop, 220,979 bp), and u35 (bottom loop, 265,136 bp) — connected by four L-records, forming an open dumbbell. B) Improved assembly graph after adding loop closure L-record (u34+ → u35−, EC = 3,576, supported by canonical-master- spanning HiFi reads in u34+ unique sequence near the wrap). The unitigs now appear as three closed loops meeting at a central junction, transforming the open dumbbell into the closed trefoil topology shown. The black segments at the central junction depict recombination repeats (37 bp u33↔u34 inverted repeat, 958 bp u33↔u35 direct repeat, 44 bp u35 self-loop, and the wrap-around edge).

Alt text: Two bandage graphs side by side depicting the mitochondrial assembly before and after adding a master-circle closure. The bandage graph on the left depicting pre-closure shows two terminal loops connected by a large single line, and the bandage graph on the left depicting post-closure exhibits a trefoil shape, together depicting multiple isomers mediated by repeat regions.

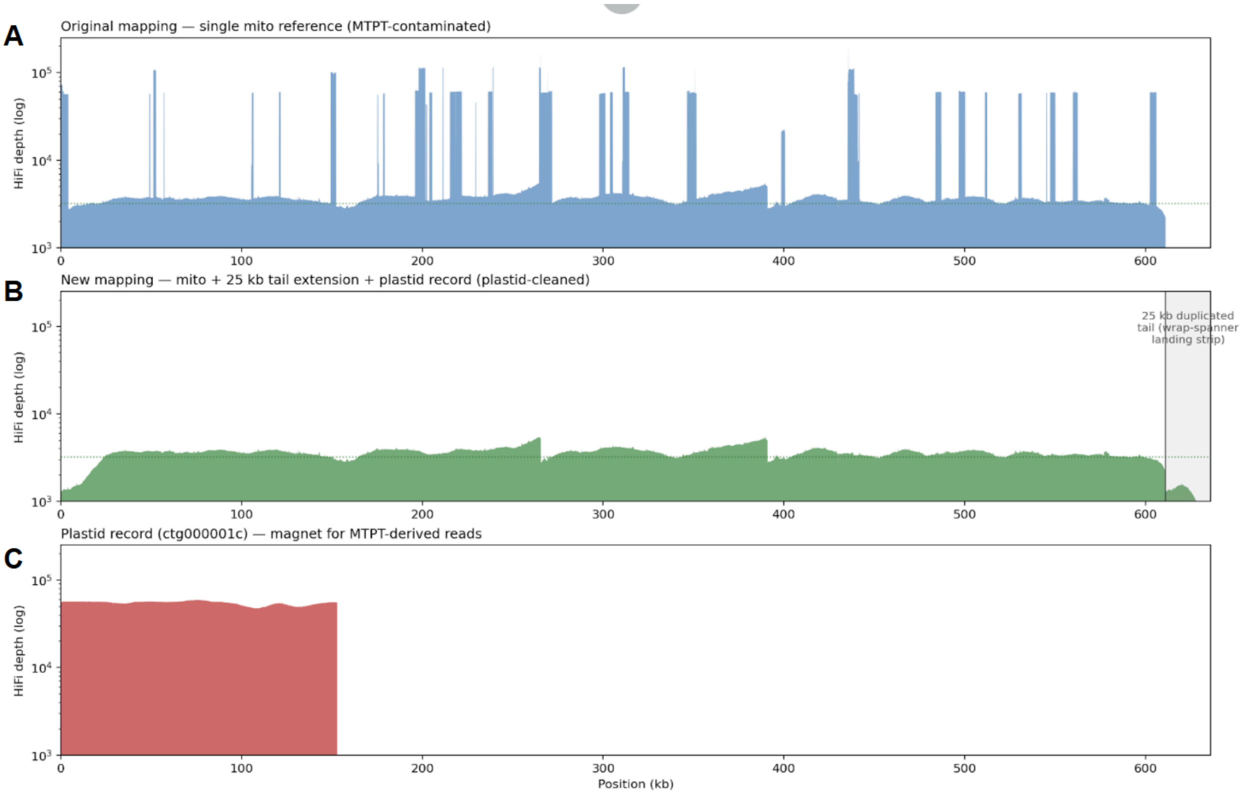


**Figure S7: Mapping coverage with and without plastid in reference.** HiFi coverage before and after using a plastid containing custom reference. A) mapping to the single-record reference: canonical mitochondrial baseline ~3,200× (dotted green) is interrupted by ≥30 MTPT-driven spikes ranging from 22,000× to 201,000×. B) new mapping to the custom two-record reference (25 kb mitochondrion appended plus plastid) with the chloroplast as a separate record: depth is uniformly ~3,000–3,600× across the entire 611 kb master with all MTPT spikes removed; the half-coverage region at the right (positions 611,107–636,107, gray-shaded) is the 25 kb duplicated tail, where wrap-spanning reads' coverage splits between the two equivalent landing strips, providing read-level confirmation of master-circle closure. C) plastid record coverage: median 55,541×, max 59,470×, consistent with the genuine plastid-genome read coverage that has been absorbed off the MTPT regions.

Alt text: A three-panel coverage plot comparing mapping with and without a plastid reference.

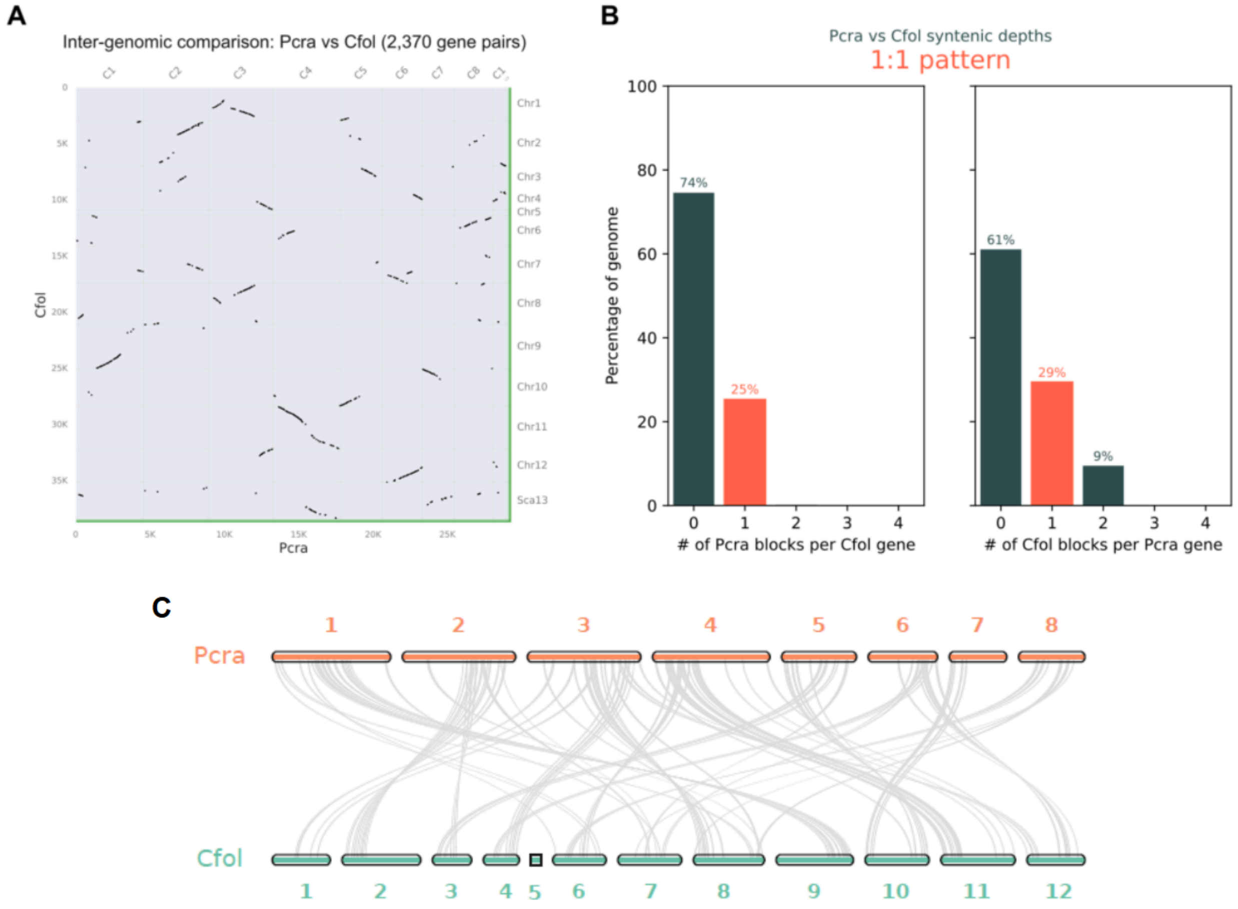


**Figure S8: Synteny between *Castilleja foliolosa* and relative *Pedicularus cranolopha*.** A) Dot plot depicting runs of synteny across the *C. foliolosa* and *P. cranolopha* genomes, denoted as Cfol and Pcra, respectively. *C. foliolosa* is depicted on the y-axis, and *P. cranolopha* is depicted on the x-axis. Each dot represents a syntenic gene pair identified via MCScanX. Diagonal lines extending from the bottom-left to top-right indicate regions of conserved gene order, while lines with negative slopes indicate chromosomal inversions. B) Bar charts illustrating the distribution of syntenic blocks per gene across both genomes, highlighting a predominantly 1:1 orthologous relationship. The left panel shows the number of Pcra blocks per Cfol gene, and the right panel shows the number of *Cfol* blocks per *Pcra* gene. C) Riparian plot depicting location of syntenic regions across *C. foliolosa* and *P. cranolopha*, depicted in blue and orange respectively, structurally scaled by their internal gene rank order generated via GENESPACE.

Alt text: Three-panel figure showing genome synteny between *C. foliolosa* and *P. cranolopha*. Panel A is a dot plot displaying conserved gene orders and inversions. Panel B uses bar charts to show a predominantly 1:1 orthologous relationship alongside large non-syntenic regions. Panel C is a ribbon-based riparian plot mapping syntenic connections between the two genomes scaled by gene rank.

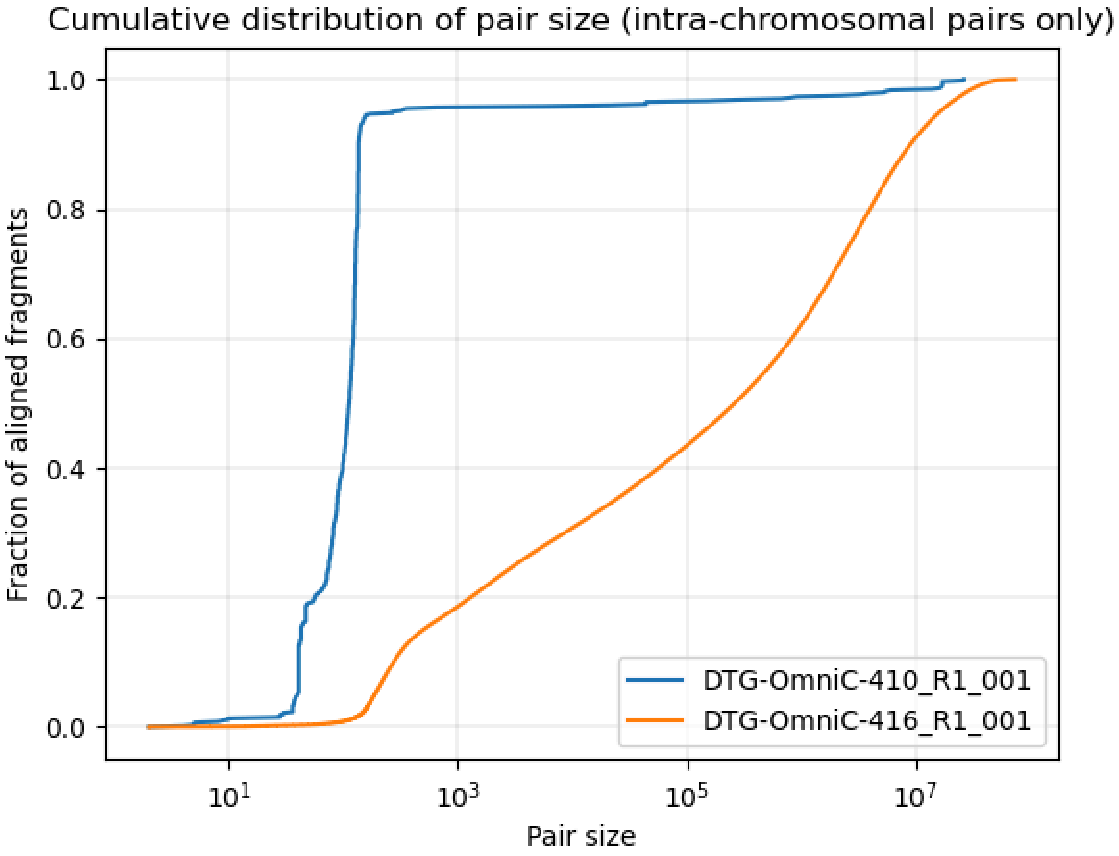


**Figure S9: Cumulative distribution of intra-chromosomal alignment pair sizes.** Fraction of aligned fragments relative to the interaction pair size in base pairs on a log scale for two sequencing libraries. The first Omni-C run (blue line) exhibits a sharp accumulation of fragments at short distances, indicating a dominance of short-range insert fragments. In contrast, the second Omni-C run (orange line) demonstrates a steady, broad cumulative increase spanning long-range distances, characteristic of successful long-range chromatin interaction capture

Alt text: Line graph showing the cumulative distribution of intra-chromosomal alignment pair sizes for two Omni-C libraries. Run 1 rises sharply near 100 base pairs and plateaus, showing mostly short-range fragments. Run 2 rises gradually across the entire scale, indicating a high proportion of long-range genomic interactions.

### Supplementary Tables:

| Class | Family | Count | Bp Masked | % Masked |
|---|---|---|---|---|
| LINE | L1 | 3452 | 4243111 | 0.83% |
| LTR | Copia | 40855 | 46843599 | 9.17% |
| | Gypsy | 96459 | 95996392 | 18.79% |
| | unknown | 177662 | 130468622 | 25.54% |
| SINE | tRNA | 264 | 60767 | 0.01% |
| TIR | CACTA | 9312 | 4164131 | 0.82% |
| | Mutator | 24968 | 8716796 | 1.71% |
| | PIF_Harbinger | 14184 | 4556622 | 0.89% |
| | Tc1_Mariner | 27676 | 6525751 | 1.28% |
| | hAT | 36390 | 11084764 | 2.17% |
| non-LTR | pararetrovirus | 135 | 99357 | 0.02% |
| nonTIR | helitron | 66254 | 21882940 | 4.28% |
| rDNA | 45S | 1089 | 558025 | 0.11% |
| Repeat fragment | Repeat fragment | 9389 | 3537545 | 0.69% |
| **TOTAL** | | **508089** | **338738422** | **66.32%** |

**Table S1: Summary of de-novo transposable element annotation statistics.** EDTA de-novo transposable element (TE) annotation divided by TE class identity, detailing the number of identified TEs, amount of the genome masked in basepairs, and the percentage of the genome masked for each TE family. Totals are summarized in bold. Abbreviations: LINE= Long Interspersed Nuclear Element, LTR= Long Terminal Repeat, SINE= Short Interspersed Nuclear Element, TIR= Terminal Inverted Repeat, and rDNA = Ribosomal DNA.

| Metric | 2024 - Burger | 2026 - Jacobs |
|---|---|---|
| **Diploid assembly length** | 752 Mbp | 784 Mbp |
| **Haploid assembly length** | 555 Mbp | 510 Mbp |
| **# of contigs** | 551 | 106 |
| **Largest contig** | 32,005,117 bp | 59,048,631 bp |
| **N50** | 11,150,047 bp | 49,877,665 bp |
| **N90** | 2,884,952 bp | 26,291,936 bp |
| **Complete BUSCOs** | 87.50% | 94.11% |
| **Gaps** | 0 | 66 |

**Table S2: Quality and contiguity statistical comparison across previously published 2024 *Castilleja foliolosa* genome and the new assembly reported in this publication.** General stats were calculated using quast and merqury. BUSCO scores for the 2024 assembly and 2026 assembly were calculated using eudicots_odb10 and embryophyta_odb10, respectively.

| Contig | Length of Reads | Number of Reads |
|---|---|---|
| tig00000002 | 114760 | 71 |
| tig00000003 | 100376 | 68 |
| tig00000004 | 40524 | 27 |
| tig00000005 | 38409 | 20 |
| tig00000006 | 28757 | 14 |
| tig00000007 | 25068 | 6 |
| tig00000008 | 27742 | 9 |
| tig00000009 | 18184 | 5 |
| tig00000010 | 28251 | 11 |
| tig00000011 | 23061 | 6 |
| tig00000012 | 53411 | 394 |
| tig00000013 | 18015 | 4 |
| tig00000014 | 17735 | 11 |
| tig00000015 | 17562 | 3 |
| tig00000016 | 20964 | 128 |
| tig00000017 | 18913 | 3 |
| tig00000018 | 121598 | 1229 |
| tig00000019 | 18200 | 10 |

**Table S3: Length and quantity of contigs for chloroplast assembly.**

| MTPT | Coordinates | Length | Peak depth | Location | Plastid gene(s) |
|---|---|---|---|---|---|
| MTPT-1 | 1–4,203 | 4,203 bp | 113,358× | u35:260,934–265,136 (-) | trnT-GGU (tRNA), psbD, psbA, psbC, psbB |
| MTPT-2 | 48,973–49,515 | 543 bp | 57,554× | u35:215,622–216,164 (-) | psbC, psbB |
| MTPT-3 | 51,538–52,708 | 1,171 bp | 107,508× | u35:212,429–213,599 (-) | rps12, rrn16 (rRNA), trnV-GAC (tRNA) |
| MTPT-4 | 56,926–57,255 | 330 bp | 58,992× | u35:207,882–208,211 (-) | cemA, petA |
| MTPT-5 | 105,814–106,597 | 784 bp | 59,840× | u35:158,540–159,323 (-) | psbJ, psbL, psbF, psbE |
| MTPT-6 | 120,773–121,618 | 846 bp | 60,147× | u35:143,519–144,364 (-) | rpoC2 |
| MTPT-7 | 149,309–152,185 | 2,877 bp | 103,415× | u35:112,952–115,828 (-) | ycf1, trnN-GUU (tRNA), trnR-ACG (tRNA), rrn5 (rRNA), rrn4.5 (rRNA), rrn23 (rRNA), rrn23-fragment (rRNA) |
| MTPT-8 | 175,136–175,555 | 420 bp | 56,806× | u35:89,582–90,001 (-) | ndhD, ndhF |
| MTPT-9 | 178,339–179,147 | 809 bp | 58,084× | u35:85,990–86,798 (-) | trnS-UGA (tRNA), psbZ |
| MTPT-10 | 196,030–202,559 | 6,530 bp | 167,162× | u35:62,578–69,107 (-) | ycf2, ndhB, trnL-CAA (tRNA), clpP1, psbB |
| MTPT-11 | 203,927–205,431 | 1,505 bp | 113,140× | u35:59,706–61,210 (-) | rps2, rpoC2 |
| MTPT-12 | 211,278–211,637 | 360 bp | 113,964× | u35:53,500–53,859 (-) | rpl2, ycf2 |
| MTPT-13 | 215,326–221,884 | 6,559 bp | 60,846× | u35:43,253–49,811 (-) | psbA, psbD, matK, rps16 |

| | | | | | |
|---|---|---|---|---|---|
| MTPT-14 | 229,444–229,644 | 201 bp | 45,383× | u35:35,493–35,693 (-) | — |
| MTPT-15 | 236,383–239,367 | 2,985 bp | 115,307× | u35:25,770–28,754 (-) | petL, petG, trnW-CCA (tRNA), trnP-UGG (tRNA), psaJ |
| MTPT-16 | 264,606–271,808 | 7,203 bp | 165,286× | u33:118,358–125,560 (-) | rpoC2, rpoC1, rpoA, rps11, rpl36, infA, rps8, rpl14 |
| MTPT-17 | 297,844–301,287 | 3,444 bp | 58,293× | u33:88,879–92,322 (-) | psaC, ndhE, ndhG, ndhI, ndhA |
| MTPT-18 | 303,894–305,348 | 1,455 bp | 60,090× | u33:84,818–86,272 (-) | rpoB, infA |
| MTPT-19 | 310,825–314,517 | 3,693 bp | 168,741× | u33:75,649–79,341 (-) | psaA, psb30, pafI, ycf2 |
| MTPT-20 | 346,381–351,681 | 5,301 bp | 112,941× | u33:38,485–43,785 (-) | psaB, psaA |
| MTPT-21 | 398,508–400,673 | 2,166 bp | 22,570× | u34:8,379–10,544 (+) | rpl14, rpl16, rps3 |
| MTPT-22 | 435,228–441,821 | 6,594 bp | 201,194× | u34:45,099–51,692 (+) | trnL-CAA (tRNA), ndhB, ndhF, rps7, rps12, ndhH, rps15, ycf1 |
| MTPT-23 | 483,872–487,115 | 3,244 bp | 60,492× | u34:93,743–96,986 (+) | ndhK, ndhC, trnM-CAU (tRNA), atpE, atpB |
| MTPT-24 | 496,717–500,260 | 3,544 bp | 115,813× | u34:106,588–110,131 (+) | trnF-GAA (tRNA), ndhJ, ndhK |
| MTPT-25 | 511,227–512,497 | 1,271 bp | 59,800× | u34:121,098–122,368 (+) | rbcL |
| MTPT-26 | 529,699–531,548 | 1,850 bp | 109,711× | u34:139,570–141,419 (+) | ndhA, rpl33, ndhH |
| MTPT-27 | 545,099–545,461 | 363 bp | 58,911× | u34:154,970–155,332 (+) | cemA |
| MTPT-28 | 547,277–550,127 | 2,851 bp | 59,617× | u34:157,148–159,998 (+) | rpoC1, rpoB |

| | | | | | |
|---|---|---|---|---|---|
| MTPT-29 | 559,880–562,401 | 2,522 bp | 115,115× | u34:169,751–172,272 (+) | psaI, rpl20 |
| MTPT-30 | 602,413–606,095 | 3,683 bp | 59,969× | u34:212,284–215,966 (+) | trnR-UCU (tRNA), atpA, atpF |

**Supplemental Table S4: MTPT Sites.** The 30 MTPT (mitochondrial–plastid transfer) regions detected in the *C. foliosa* mitochondrial master path, with concat-path coordinates, length, apparent peak HiFi coverage, original-unitig location and orientation, and the chloroplast gene(s) overlapped by the homologous plastid alignment(s). Peak depths are samtools depth maxima from the full read mappings reflecting contamination from chloroplast-derived HiFi reads that co-map to MTPT regions; true mito-only depths are recovered after plastid-magnet partitioning (see Methods). Feature-type qualifiers (tRNA) and (rRNA) are shown; unqualified names are protein-coding (CDS) genes. Plastid loci are from our GeSeq annotation of the *C. foliosa* chloroplast assembly.

| Isoform | Walk | Approx. size |
|---|---|---|
| 1 (master, canonical) | u35+ → u33+ → u34± → u35+ | 612 kb |
| 2 (subgenomic) | u35+ → u35+ | 265 kb |

**Table S5: Isoforms enumerated from the GFA.** Sizes are the sum of segment lengths in the walk and do not subtract overlap regions. Isoform 1 is the canonical master circle.

| Isoform | Estimated abundance |
|---|---|
| Master, canonical | ~99.7% |
| Master, recombinant A | ~0.10% |
| Master, recombinant B | ~0.05% |
| u35 subgenomic circle | ~0.1% |

**Table S6: Estimated relative abundance of each mitochondrial structural isoform.** The three sub-stoichiometric variants are at comparable, low abundance, consistent with infrequent intermolecular recombination at imperfect or short repeats.

| Species | Genome Size | BioProject | BioSample | Source | Link |
|---|---|---|---|---|---|
| Lindenbergia sp. Thulin 8079 | 242.17 Gb | PRJCA008847 | SAMC695115 | Xu et al. (2021) | https://ngdc.cncb.ac.cn/gwh/Assembly/24466/show |
| Phelipanche aegyptiaca | 3.88 Gb | PRJCA008847 | SAMC695117 | Xu et al. (2021) | https://ngdc.cncb.ac.cn/gwh/Assembly/24467/show |
| Orobanche cernua var. cumana | 1.46 Gb | PRJCA008847 | SAMC695116 | Xu et al. (2021) | https://ngdc.cncb.ac.cn/gwh/Assembly/24468/show |
| Coffea canephora | 568.6 Mb | PRJEB4211 | SAMEA3146290 | Denoeud, et al. (2014). | https://www.ncbi.nlm.nih.gov/datasets/genome/GCA_900059795.1/ |
| Olea Europea | 1.1 Gb | PRJNA350614 | SAMN05943011 | Unver, et al. (2017) | https://www.ncbi.nlm.nih.gov/datasets/genome/GCF_002742605.1/ |
| Erythranthe guttata | 321.6 Mb | PRJNA13880 | SAMN02742818 | Hellsten, et al. (2013) | https://www.ncbi.nlm.nih.gov/datasets/genome/GCF_000504015.1/ |
| Striga asiatica | 471.6 Mb | PRJDB4996 | SAMD00056127 | Yoshida, et al. (2019) | https://www.ncbi.nlm.nih.gov/datasets/genome/GCA_008636005.1/ |
| Phtheirospermum japonicum | 1.2 Gb | PRJDB3858 | SAMD00029051 | in prep | https://www.ncbi.nlm.nih.gov/datasets/genome/GCA_014905375.1/ |
| Pedicularis cranolopha | NA | NA | NA | in prep | NA |

**Table S7: Location of genomic resources used in comparative genomics.** *Pedicularis cranolopha* sequence was obtained from the Eaton Lab at Columbia University.

171

| GO Term | Ontology ID | Number of *Castilleja*-specific Genes | Number of Reference Genes | P-val |
|---|---|---|---|---|
| methylenetetrahydrofolate reductase [NAD(P)H] activity | GO:0004489 | 4 | 4 | 4.76E-41 |
| L-methionine metabolic process | GO:0006555 | 4 | 4 | 4.76E-41 |
| nucleoside transmembrane transporter activity | GO:0005337 | 4 | 5 | 6.81E-33 |
| nucleoside transmembrane transport | GO:1901642 | 4 | 5 | 6.81E-33 |
| zinc ion binding | GO:0008270 | 79 | 1237 | 2.30E-25 |
| DNA replication-dependent nucleosome assembly | GO:0000723 | 5 | 10 | 2.55E-25 |
| DNA-directed DNA polymerase activity | GO:0015074 | 2 | 2 | 1.69E-21 |
| peptide transport | GO:0015833 | 2 | 2 | 1.69E-21 |
| peptide transmembrane transporter activity | GO:1904680 | 2 | 2 | 1.69E-21 |
| ceramide biosynthetic process | GO:0046513 | 5 | 12 | 4.62E-21 |
| demethylase activity | GO:0050291 | 5 | 12 | 4.62E-21 |
| transcription termination | GO:0006353 | 3 | 6 | 6.19E-16 |

| | | | | |
|---|---|---|---|---|
| diphosphate-fructose-6-phosphate 1-phosphotransferase activity | GO:0047334 | 3 | 6 | 6.19E-16 |
| mitochondrion organization | GO:0007005 | 3 | 7 | 9.91E-14 |
| nucleic acid binding | GO:0003676 | 60 | 1171 | 1.09E-12 |
| heat shock protein binding | GO:0031072 | 5 | 21 | 6.49E-12 |
| ribosome biogenesis | GO:0000350 | 1 | 1 | 1.20E-11 |
| chromosome, telomeric region | GO:0000781 | 1 | 1 | 1.20E-11 |
| primary amine oxidase activity | GO:0003852 | 1 | 1 | 1.20E-11 |
| IMP cyclohydrolase activity | GO:0003937 | 1 | 1 | 1.20E-11 |
| formate-tetrahydrofolate ligase activity | GO:0004329 | 1 | 1 | 1.20E-11 |
| hydroxyacylglutathione hydrolase activity | GO:0004416 | 1 | 1 | 1.20E-11 |
| monophenol monooxygenase activity | GO:0004514 | 1 | 1 | 1.20E-11 |
| phosphoglycerate mutase activity | GO:0004643 | 1 | 1 | 1.20E-11 |
| tubulin complex assembly | GO:0007021 | 1 | 1 | 1.20E-11 |
| intracellular transport | GO:0007023 | 1 | 1 | 1.20E-11 |
| leucine biosynthetic process | GO:0009098 | 1 | 1 | 1.20E-11 |
| methylglyoxal catabolic process | GO:0019243 | 1 | 1 | 1.20E-11 |
| plasma membrane respiratory chain | GO:0045273 | 1 | 1 | 1.20E-11 |

| | | | | |
|---|---|---|---|---|
| vesicle fusion | GO:0048280 | 1 | 1 | 1.20E-11 |
| synaptic vesicle endocytosis | GO:0048487 | 1 | 1 | 1.20E-11 |
| neurotransmitter transport | GO:0042138 | 2 | 4 | 3.28E-11 |
| chromatin assembly | GO:0006338 | 4 | 16 | 2.30E-10 |
| beta-glucuronidase activity | GO:0004563 | 2 | 5 | 3.91E-09 |
| beta-xylosidase activity | GO:0008184 | 2 | 5 | 3.91E-09 |
| DNA helicase activity | GO:0003678 | 4 | 19 | 9.85E-09 |
| solute:proton symporter activity | GO:0015297 | 13 | 147 | 1.67E-08 |
| ribonuclease H activity | GO:0004527 | 3 | 14 | 4.50E-07 |
| xenobiotic transmembrane transporter activity | GO:0042910 | 9 | 96 | 7.50E-07 |
| transporter activity | GO:0005215 | 6 | 49 | 7.71E-07 |
| mitochondrial transport | GO:0006850 | 2 | 7 | 9.61E-07 |
| thylakoid | GO:0009579 | 2 | 7 | 9.61E-07 |
| phosphorelay sensor kinase activity | GO:0000151 | 1 | 2 | 2.00E-06 |
| pantothenate kinase activity | GO:0004594 | 1 | 2 | 2.00E-06 |
| phosphomethylpyrimidine kinase activity | GO:0004649 | 1 | 2 | 2.00E-06 |
| ubiquitin recognition component activity | GO:0034450 | 1 | 2 | 2.00E-06 |

| | | | | |
|---|---|---|---|---|
| protein transport | GO:0015035 | 2 | 8 | 5.44E-06 |
| mitochondrial outer membrane | GO:0005741 | 3 | 17 | 6.95E-06 |
| 6-phosphofructokinase activity | GO:0003872 | 3 | 18 | 1.42E-05 |
| DNA topoisomerase (ATP-hydrolyzing) activity | GO:0003918 | 2 | 9 | 2.11E-05 |
| sucrose transmembrane transporter activity | GO:0005384 | 2 | 9 | 2.11E-05 |
| structural constituent of muscle | GO:0030026 | 2 | 9 | 2.11E-05 |
| calcium ion binding | GO:0005509 | 19 | 369 | 4.84E-05 |
| chorismate mutase activity | GO:0004106 | 1 | 3 | 0.000121 |
| fructose-bisphosphate aldolase activity | GO:0004348 | 1 | 3 | 0.000121 |
| base-excision repair | GO:0006282 | 1 | 3 | 0.000121 |
| glucosylceramide biosynthetic process | GO:0006680 | 1 | 3 | 0.000121 |
| coenzyme A biosynthetic process | GO:0015937 | 1 | 3 | 0.000121 |
| vitamin D receptor binding | GO:0042819 | 1 | 3 | 0.000121 |
| vitamin L receptor binding | GO:0042823 | 1 | 3 | 0.000121 |
| glutamate synthase (NADH) activity | GO:0046417 | 1 | 3 | 0.000121 |
| ubiquitin ligase binding | GO:0043130 | 4 | 37 | 0.00018 |
| extracellular region | GO:0005576 | 5 | 55 | 0.000249 |

| | | | | |
|---|---|---|---|---|
| unfolded protein binding | GO:0051082 | 5 | 56 | 0.000302 |
| iron ion transport | GO:0006820 | 2 | 12 | 0.000328 |
| phosphoinositide binding | GO:0035091 | 4 | 40 | 0.000394 |
| phospholipid binding | GO:0005543 | 3 | 26 | 0.000606 |
| DNA metabolic process | GO:0006259 | 2 | 13 | 0.000621 |
| antioxidant activity | GO:0016209 | 2 | 13 | 0.000621 |
| structural constituent of ribosome | GO:0003735 | 16 | 339 | 0.000735 |
| 2-isoprenyl diphosphate binding | GO:0005545 | 3 | 27 | 0.000831 |
| clathrin-coated vesicle | GO:0030136 | 3 | 27 | 0.000831 |
| clathrin coat cloaking | GO:0048268 | 3 | 27 | 0.000831 |
| purine nucleotide biosynthetic process | GO:0006164 | 1 | 4 | 0.000993 |
| cytoduction | GO:0008373 | 1 | 4 | 0.000993 |
| magnesium ion transmembrane transporter activity | GO:0015098 | 1 | 4 | 0.000993 |
| copper ion transport | GO:0015689 | 1 | 4 | 0.000993 |
| intramolecular transferase activity | GO:0016866 | 1 | 4 | 0.000993 |
| amino acid import across plasma membrane | GO:0070652 | 1 | 4 | 0.000993 |
| plant-type vacuole membrane | GO:0071669 | 1 | 4 | 0.000993 |
| translation | GO:0006412 | 16 | 350 | 0.001186 |

| | | | | |
|---|---|---|---|---|
| serine-type peptidase activity | GO:0008236 | 9 | 161 | 0.001703 |
| fructose metabolic process | GO:0006002 | 2 | 15 | 0.001746 |
| clathrin binding | GO:0030276 | 3 | 31 | 0.002411 |
| nucleus | GO:0005634 | 3 | 592 | 0.002554 |
| pyruvate kinase activity | GO:0004751 | 1 | 5 | 0.003615 |
| galactokinase activity | GO:0009052 | 1 | 5 | 0.003615 |
| transferase activity, transferring pentosyl groups | GO:0016763 | 1 | 5 | 0.003615 |
| extracellular space | GO:0019898 | 2 | 17 | 0.003881 |
| L-ascorbic acid binding | GO:0031418 | 2 | 17 | 0.003881 |
| cation transport | GO:0006812 | 5 | 74 | 0.004004 |
| protein folding | GO:0006457 | 5 | 75 | 0.004458 |
| protein ubiquitination | GO:0016567 | 7 | 124 | 0.0047 |
| progesterone receptor binding | GO:0044772 | 2 | 18 | 0.005432 |
| ribosome | GO:0005840 | 14 | 329 | 0.005605 |
| transcription corepressor activity | GO:0003714 | 1 | 6 | 0.008742 |
| NAD biosynthetic process | GO:0009435 | 1 | 6 | 0.008742 |
| peroxisome fission | GO:0016559 | 1 | 6 | 0.008742 |
| ribonuclease T2 activity | GO:0033897 | 1 | 6 | 0.008742 |
| spindle midzone assembly | GO:0051225 | 1 | 6 | 0.008742 |

| | | | | |
|---|---|---|---|---|
| hydrolase activity, hydrolyzing O-glycosyl compounds | GO:0004553 | 14 | 348 | 0.010657 |
| hydrolase activity | GO:0016787 | 14 | 349 | 0.010999 |
| photosynthesis | GO:0015979 | 4 | 61 | 0.011237 |
| systemic acquired resistance | GO:0009654 | 2 | 21 | 0.012412 |
| protein peptidyl-prolyl isomerase activity | GO:0000413 | 3 | 41 | 0.014304 |
| cyclin-dependent protein kinase regulator activity | GO:0016538 | 2 | 22 | 0.015599 |
| serine-type endopeptidase activity | GO:0004252 | 9 | 202 | 0.015755 |
| mitochondrial proton-transporting ATP synthase complex | GO:0000276 | 1 | 7 | 0.016698 |
| copper ion transmembrane transporter activity | GO:0005375 | 1 | 7 | 0.016698 |
| metallopeptidase activity | GO:0030570 | 1 | 7 | 0.016698 |
| copper ion import across plasma membrane | GO:0035434 | 1 | 7 | 0.016698 |
| plant cell wall organization | GO:0009664 | 3 | 43 | 0.018568 |
| peroxisomal membrane | GO:0005778 | 1 | 8 | 0.027491 |
| outer membrane | GO:0019867 | 1 | 8 | 0.027491 |
| sequence-specific DNA binding | GO:0043565 | 1 | 250 | 0.03105 |
| chromosome | GO:0005694 | 2 | 26 | 0.033063 |

| | | | | |
|---|---|---|---|---|
| ubiquitin protein ligase activity | GO:0061630 | 2 | 26 | 0.033063 |
| ubiquitin-protein transferase activity | GO:0004842 | 7 | 160 | 0.035013 |
| biquitin-protein ligase inhibitor activity | GO:0030527 | 3 | 49 | 0.036023 |
| peroxisome organization | GO:0070042 | 1 | 9 | 0.040963 |
| metal ion transport | GO:0030001 | 3 | 51 | 0.04351 |
| metal ion binding | GO:0046873 | 3 | 51 | 0.04351 |
| ADP binding | GO:0043531 | 2 | 298 | 0.043626 |
| metal ion binding | GO:0046872 | 5 | 493 | 0.043688 |
| mitochondrial inner membrane | GO:0005743 | 2 | 28 | 0.044735 |
| proteolysis | GO:0006508 | 17 | 512 | 0.047566 |

**Table S8: Gene Ontological term enrichment for *Castilleja*-specific genes**. Gene Ontology (GO) term enrichment was calculated by transferring GO terms from the closely related *Mimulus guttatus* (IM62 v3.1; Lovell et al., 2025) reference genome using BLASTP, retaining the top-scoring hit per query above an e-value threshold of ≤ 1e-05. GO annotations were retrieved using Phytozome’s Biomart Tool (Goodstein et al., 2012). Enrichment was performed using Planteome’s Ontology Enrichment Analysis Tool (https://planteome.org/oat/), applying a chi-squared test. Terms are in descending order by significance.
